# Gut microbiota has a widespread and modifiable effect on host gene regulation

**DOI:** 10.1101/210294

**Authors:** Allison L Richards, Amanda L Muehlbauer, Adnan Alazizi, Michael B Burns, Anthony Findley, Francesco Messina, Trevor J Gould, Camilla Cascardo, Roger Pique-Regi, Ran Blekhman, Francesca Luca

## Abstract

Variation in gut microbiome is associated with wellness and disease in humans, yet the molecular mechanisms by which this variation affects the host are not well understood. A likely mechanism is through changing gene regulation in interfacing host epithelial cells. Here, we treated colonic epithelial cells with live microbiota from five healthy individuals and quantified induced changes in transcriptional regulation and chromatin accessibility in host cells. We identified over 5,000 host genes that change expression, including 588 distinct associations between specific taxa and host genes. The taxa with the strongest influence on gene expression alter the response of genes associated with complex traits. Using ATAC-seq, we show that a subset of these changes in gene expression are likely the result of changes in host chromatin accessibility and transcription factor binding induced by exposure to gut microbiota. We then created a manipulated microbial community with titrated doses of *Collinsella*, demonstrating that both natural and controlled microbiome composition leads to distinct, and predictable, gene expression profiles in host cells. Together, our results suggest that specific microbes play an important role in regulating expression of individual host genes involved in human complex traits. The ability to fine tune the expression of host genes by manipulating the microbiome suggests future therapeutic routes.

## Introduction

The microbial community in the human digestive tract, the gut microbiota, is highly complex, and also displays strong variation across individuals [22, 32, 43]. Variability in gut microbiota composition is related to many factors, including medication, diet and genetics [4, 5, 13, 15, 23, 30, 33, 35, 49, 53, 55, 56, 58]. The gut microbiota has a variety of functions within the host, such as metabolism of certain compounds [1, 18, 21, 46], and its composition is correlated with several diseases, such as Crohn’s disease and colorectal cancer [7, 19, 29, 41, 47, 50]. In mice, certain microbial communities can lead to changes in the host’s weight and overall health, suggesting that there is a reciprocal effect between the host and the gut microbiota [20]. Recent work in mice has explored host gene expression and gene regulation in response to microbiome exposure [8, 11, 16]. These studies suggested that the microbiome can induce both epigenetic changes and binding of specific transcription factors [8, 11, 28, 44, 48]. However, in humans, our ability to study the effects and mechanism of the microbiome *in vivo* is severely limited. Recently, we have described an in vitro approach based on human epithelial cells inoculated with live microbial communities [45] that is well suited to study the effects of the microbiome on human gene regulation. Here we seek to use this in vitro system to determine the extent and mechanism by which inter-individual variation in microbiome composition drives differences in gene expression in the host cells. We also seek to determine if specific microbial taxa drive gene expression variation, and if these changes are predictable, i.e. they can be recapitulated by manipulating the composition of the microbiome. These open questions are crucial for understanding the causal role of the microbiome in host physiology and designing targeted therapies revolving around interventions on the gut microbiome.

## Results

### Exposure to Microbiota Influences Host Gene Expression

To determine the impact of variation in the gut microbiota on host cells, we treated human colonic epithelial cells with live gut microbiota extracted from 5 healthy, unrelated, human individuals (Fig 1A, Fig S1 and Table S1). The composition of these samples is representative of other healthy gut microbiome samples from the Human Microbiome Project (Fig S2) [22, 36]. We then separately assessed changes in gene expression and microbial composition following 1, 2 and 4 hours of exposure. The overall changes in gene expression between each microbiome treatment and control cluster first by time-point (Fig 1B, Table S2) where the strongest response occurs at 2 hours following exposure (3,240 genes across any of the five microbiota samples, BH FDR < 10%, |log_2_ FC|> 0.25). Among these, we identified 669 transcripts (188 genes) that are differentially expressed in all five treatments following 2 hours of treatment (s (Fig 1C, 1 and 4 hour comparisons in Figs S3A and B). To identify genes that change consistently across the treatments at each time point, we removed the individual effect from the model and considered the 5 microbiota treatments to be replicates. Fig 1D shows two representative genes, *PDLIM5* and *DSE*, that are found to be differentially expressed at each time point. Notably, these genes show that we are able to identify various expression patterns through this model as long as the 5 microbiota treatments lead to the same response at a given time point. (Figs S4A and B, Table S3). These 5,413 genes with shared expression changes, are enriched for genes that function in protein translation, as well as those on the cell surface, such as in adherens junctions (BH FDR < 10^−6^) (Table S4, Fig S5), suggesting a biological function for consistent changes in gene expression that may relate to the host cell’s interaction with the microbiota.

**Figure 1:**
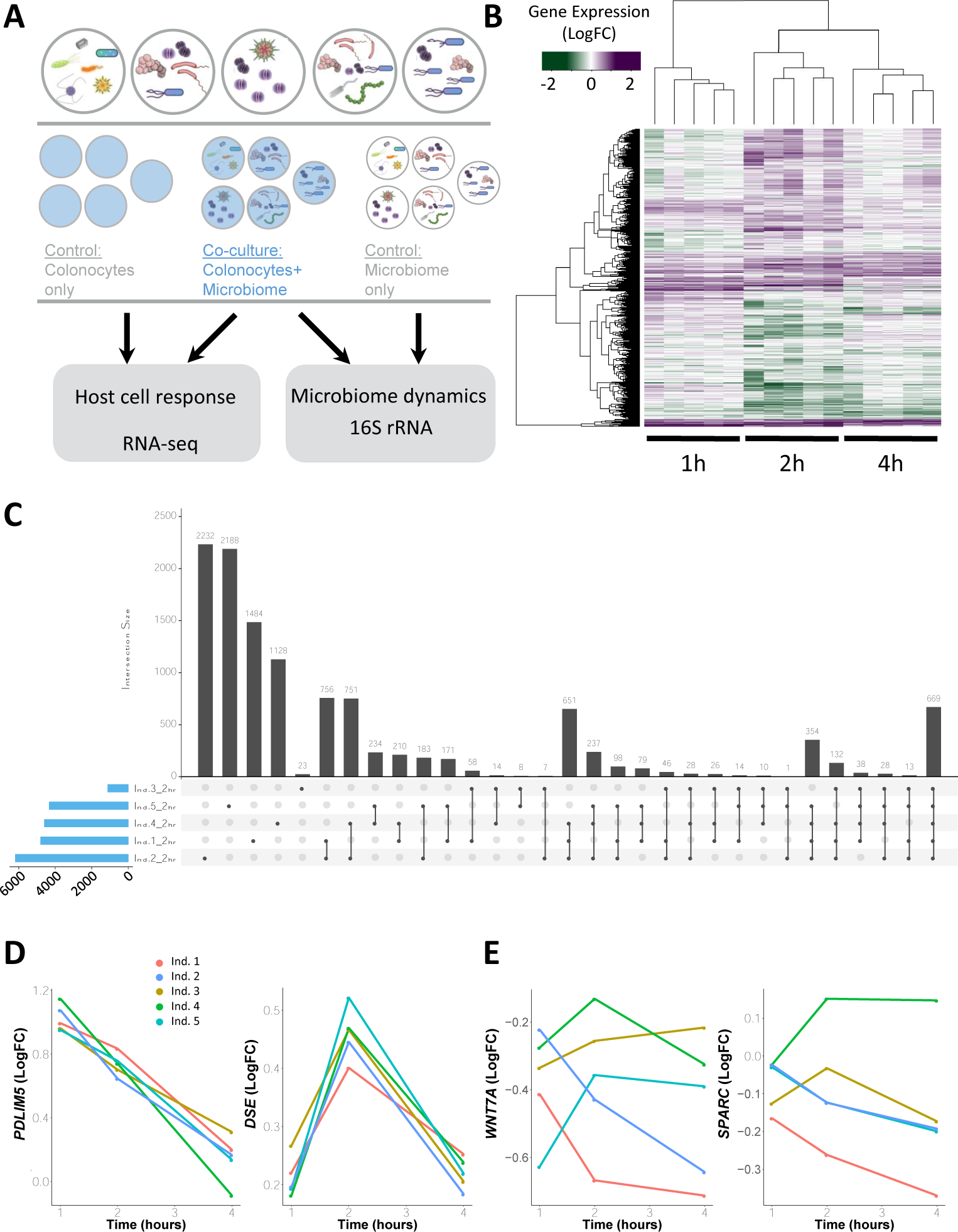
Gene expression changes in Colonocytes Treated with Microbiota from Five Unrelated Individuals. **A** Study design. Human colonocytes were inoculated separately with five microbiota samples from unrelated individuals. **B** Heatmap of gene expression changes induced at each time point by the individual microbiota samples. Purple denotes an increase in gene expression (green shows a reduction) compared to the gene expression in the control (colonocytes cultured alone). Only genes that are differentially expressed in at least one sample are shown. **C** Comparison of transcripts differentially expressed at 2 hours across the five treatments. The blue bars to the left show the total number of differentially expressed transcripts in the given set. The gray vertical bars show the number of transcripts that are in the set denoted below them. Sets with a single dark gray circle show the number of differentially expressed transcripts unique to that sample. **D** Examples of genes (*PDLIM5* and *DSE*) whose changes in expression are consistent across treatments with the five different microbiota. Changes in expression (y-axis) are shown as log2 fold change as compared to control. **E** Examples of genes (*WNT7A* and *SPARC*) whose changes in expression are significantly different across treatments with the five microbiota samples.

Each microbiota sample is derived from a different individual with a unique diet and genetic makeup. Therefore, we expect that the microbial composition and diversity of each sample differs. When we considered the uncultured microbiome, we found variability in their microbial composition and diversity (Simpson’s index range between 0.94 to 0.98). In culture, we found that microbial communities do not change dramatically over time and the microbiome maintains individual-specific composition during culture (Fig S1). When we next considered how the human colonocytes influenced microbial composition, we found that most taxa were unaffected by the presence of human cells, while 13 taxa showed varying abundance dependent on the presence of host cells (likelihood ratio test, BH FDR < 10%, examples in Fig S6, Table S5). In order to determine how the microbiota composition of each sample influences host gene expression differently, we utilized a likelihood ratio test to compare models including or excluding the individual microbiota effect. This test is able to incorporate gene expression changes over time and compare the trajectory of expression in response to the 5 different microbiota treatments. In this way, we identified 409 genes (1,484 transcripts, BH FDR < 10%, Table S6) with expression patterns that significantly differ in response to the five microbiota samples. Two examples of representative genes with differing gene expression patterns are *WNT7A* and *SPARC* (Fig 1E). These genes demonstrate that no two microbiota samples induce the same response in the genes identified by the likelihood ratio test and further show that genes have different responses to the same treatment. These data demonstrate that both the host and the microbiota influence each other and that inter-individual variation in the microbiome can lead to different gene expression responses in interacting host cells.

### Specific Microbes Influence Unique Host Genes

We hypothesized that the differences in gene expression response to each microbiome could be attributed to specific microbiota features, such as the abundance of specific taxa. For this reason we used DESeq2 to study the association between host gene expression (147,555 transcripts) and the abundance of microbial taxa (62 taxa that pass filtering criteria; see Methods and Materials) at the time of treatment. Across all possible associations (9,125,927 tests) we identified 588 transcript-by-taxon pairs with a significant association (BH FDR = 10%, Table S7), corresponding to 121 host genes with changes in expression associated with the abundance of 46 taxa. 35 of these taxa were associated with the expression of more than one host gene (BH FDR = 10%) suggesting that a single microbe may affect the regulation of many genes, and suggesting a trans regulatory mechanism by which microbes may influence host organism traits. Of the 121 host genes whose expression is associated with abundance of microbial taxa, 70 genes (219 transcripts) were also differentially expressed when each microbiota treatment was compared to control conditions, and formed two clusters with distinct functions (Fig 2A and D, examples in Fig 2B and C). Genes in the first cluster are positively correlated with genera *Ruminococcus*, *Coprococcus* and *Streptococcus*, and have functions in cell junction assembly (BH FDR < 10^−5^, Fig 2E), while the second cluster of genes, positively correlated with microbial genera including *Odoribacter*, *Blautia* and *Collinsella*, function in protein targeting to the endoplasmic reticulum (BH FDR < 10^−7^, Fig 2F). When we considered the abundances of the microbes in each cluster, we found them highly correlated, further suggesting that the microbial community as a whole may act on host gene expression (Fig S7).

**Figure 2:**
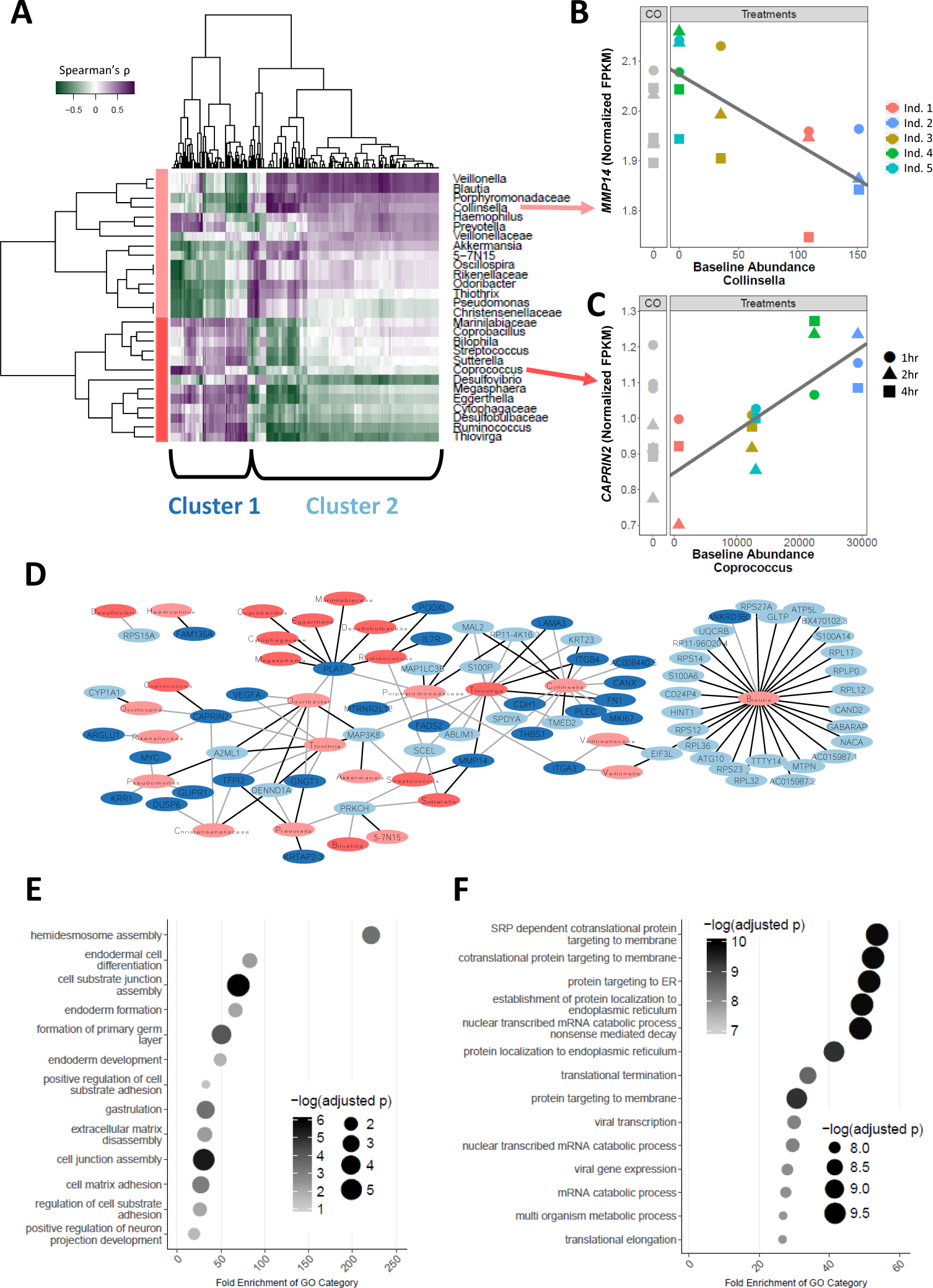
Abundance of microbiome taxa is associated with specific host gene expression changes. **A** Heatmap of microbiota taxa and colonocyte gene expression correlation (Spearman’s ρ). Rows correspond to 28 microbiota taxa and columns correspond to 219 transcripts (70 genes) that had at least one significant association (LRT). Taxa and transcripts are each clustered via hierarchical clustering showing two major groups indicated by a different shade of red (taxa) / blue (transcripts). **B and C** Examples (*MMP14*, and *CAPRIN2*) of significant association (BH FDR = 7% for both genes) between host gene expression (FPKM quantile normalized) and baseline abundance of specific taxa. **D** Network of associations between taxa and genes from the heatmap. Nodes in blue denote genes while nodes in red denote microbial taxa. Color shading indicates clusters of genes or taxa as defined in the heatmap. Black edges indicate a positive correlation while light gray indicates a negative correlation. **E and F** Gene ontology enrichment for cluster 1 and 2 respectively.

We then focused on microbial taxa associated with changes in expression for a large number of genes, since these microbes are more likely to impact host traits. To test this hypothesis, we focused on the six microbial genera that were associated with the largest number of host genes (at least 30 host genes at *p*-value < 3.5 × 10^−5^): *Odoribacter*, *Streptococcus*, *Blautia*, *Thiovirga*, *Thiothrix*, and *Collinsella*. Indeed, all taxa, except for *Blautia*, led to expression changes in genes that are 2.7-fold enriched for complex traits (*p*-value < 0.005, Fig 3A, Table S8 and Table S9) [57]. Moreover, we identified 21 genes that were associated with traits already linked to the gut microbiome, including colorectal cancer [7, 50], obesity [20, 54], and Inflammatory Bowel Disorder (IBD) [3, 17, 27]. Half of the genes (15/30) associated with the genus *Collinsella* are 3.4-fold enriched for association with a trait in GWAS (*p*-value = 0.001, Fig 3A). Previous studies have found that the abundance of *Collinsella* is correlated with several diseases, including colorectal cancer [37], Type 2 Diabetes [9] and irritable bowel syndrome [25]. Interestingly, we identified a gene, *GLTP*, that is involved in glycolipid transfer and has been associated with metabolic syndrome [59], whose expression is influenced by the abundance of the genus *Collinsella* in each of the five microbiota samples (BH FDR = 12.6%). This suggests that microbes of the genus *Collinsella* may influence metabolic syndrome in the host through regulation of genes in the colon, such as *GLTP*. These data also suggest that specific microorganisms, and not simply general exposure to the entire gut microbiota, can lead to changes in many genes’ expression. Furthermore, these results support the hypothesis that variation in the abundance of members of the microbiota may influence complex traits.

**Figure 3:**
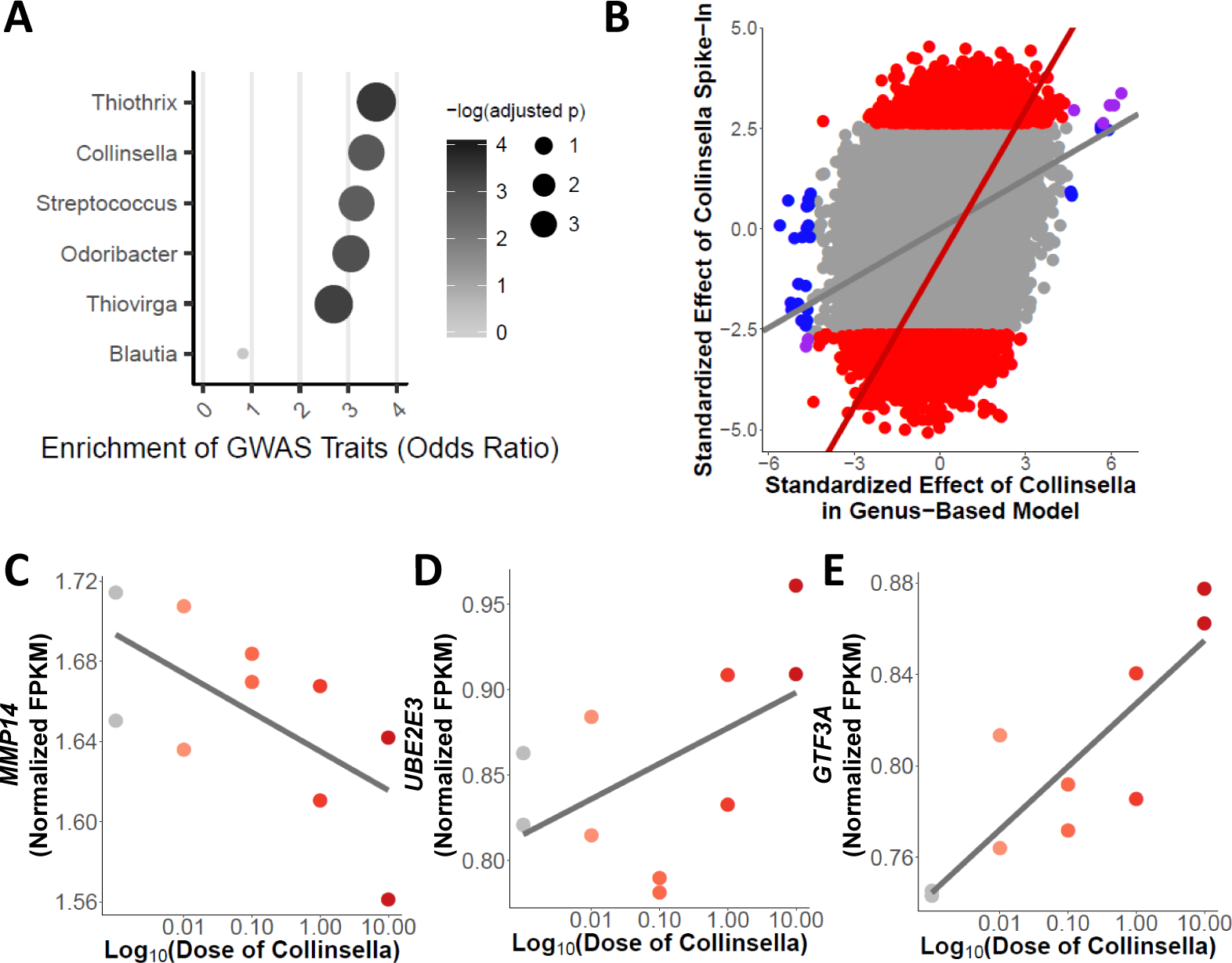
Manipulation of the Microbial Community Induces Predictable Gene Expression Changes in the Host. **A** Enrichment for complex traits across genes with changes in expression associated with microbial taxa (BH FDR < 20%). **B** Scatterplot of the effect of *Collinsella* abundance from the five microbiome samples (x-axis) and effect of *Collinsella* aerofaciens from the spike-in validation experiment (y-axis). Plotted are log_2_ fold changes normalized by the standard error. Blue points denote transcripts that are DE only in the genus-based model including abundance of *Collinsella*. The red points and line highlight the transcripts that are DE in the spike-in experiment. Gray points denote transcripts that are not differentially expressed in either the gene-based model with abundance of *Collinsella* or in the spike-in experiment. There is a correlation across all points (p-value < 10^−20^, *ρ*= 0:29) and across transcripts differentially expressed in the spike-in experiment (p-value < 10^−20^, *ρ* = 0:46). **C-E** Examples (*MMP14*, *UBE2E3*, and *GTF3A*) of significant association (DESeq2 BH FDR = 9%, 6% and 7%, respectively) between host gene expression (FPKM quantile normalized) and abundance of Collinsella aerofaciens from spike-in validation experiment (linear regression *p*-value = 0:05, 0:02, and 0:004 with Pearson’s *r* = −0:64, 0:73, and 0:81, respectively).

### Validation of Changes in Host Gene Expression Due to Specific Microbiota

To validate and further demonstrate the effect of specific microbes on host gene expression, we treated colonocytes with a microbiota sample without any detectable *Collinsella aerofaciens*, and supplemented it with titrated abundances of this bacterium relative to the whole microbiota sample: 0.01%, 0.1%, 1% and 10%. We used RNA-sequencing to study the resulting changes in gene expression, and identified 1,570 genes that change expression (BH FDR = 10%, Table S10 and Fig S8) depending on the abundance of *Collinsella aerofaciens*. When we consider the changes in gene expression associated with *Collinsella* abundance in the five microbiota treatments, we found that the effects of *Collinsella* in both experiments are correlated (Fig 3B). We validate 19 out of 29 genes (*p*-value = 0.0002, OR = 4.1), originally identified (BH FDR = 20%), including *GLTP* and *MMP14* (original BH FDR = 7% in Fig 2B, spike-in validation BH FDR = 9% in Fig 3C), demonstrating that *Collinsella* is sufficient for changes in the expression of these genes. The large number of genes that change expression in this experiment could be due to several factors, including the increase in power from a larger number of samples. These 1,570 genes are enriched for genes associated with complex traits from GWAS (*p*-value = 10^−10^, OR = 1.5) and specifically enriched for genes associated with HDL cholesterol (Bonferroni corrected *p*-value = 0.018, OR = 2.75). Other DE genes are associated with additional traits relevant to the microbiome, such as body mass index (like *GTF3A* in Fig 3E), obesity and Celiac disease (like *UBE2E3* in Fig 3D). This spike-in experiment shows that host gene expression can be modulated by changing the abundance of a single bacterial species within the microbiome.

### Changes in Chromatin Accessibility and Transcription Factor Binding Following Microbiota Exposure

In order to investigate the regulatory mechanism whereby the microbiome induces changes in host gene expression, we performed ATAC-seq in colonocytes inoculated for two hours with each of the five microbiota samples and were able to identify ATAC-seq peaks at the TSS of expressed genes (Fig S13). We then used DESeq2 to characterize regions of differential chromatin accessibility. We identified only a limited number of regions differentially accessible across the five microbiome treatments with two technical replicates (234, within 50kb of genes, Fig S9 and Fig S10). Nevertheless, we found an enrichment for differentially accessible regions at 2 hours within 50kb of genes differentially expressed at 4 hours (Fig 4A and Table S11, *p*-value = 3.96 × 10^−4^, OR = 2.13). Interestingly, we did not find the same enrichment when we examined genes differentially expressed at 1 and 2 hours following exposure to gut microbiota (*p*-value > 0.15), suggesting that the changes in chromatin accessibility that we identified occur first (at 2 hours), and then lead to subsequent changes in gene expression by 4 hours post-inoculation.

**Figure 4:**
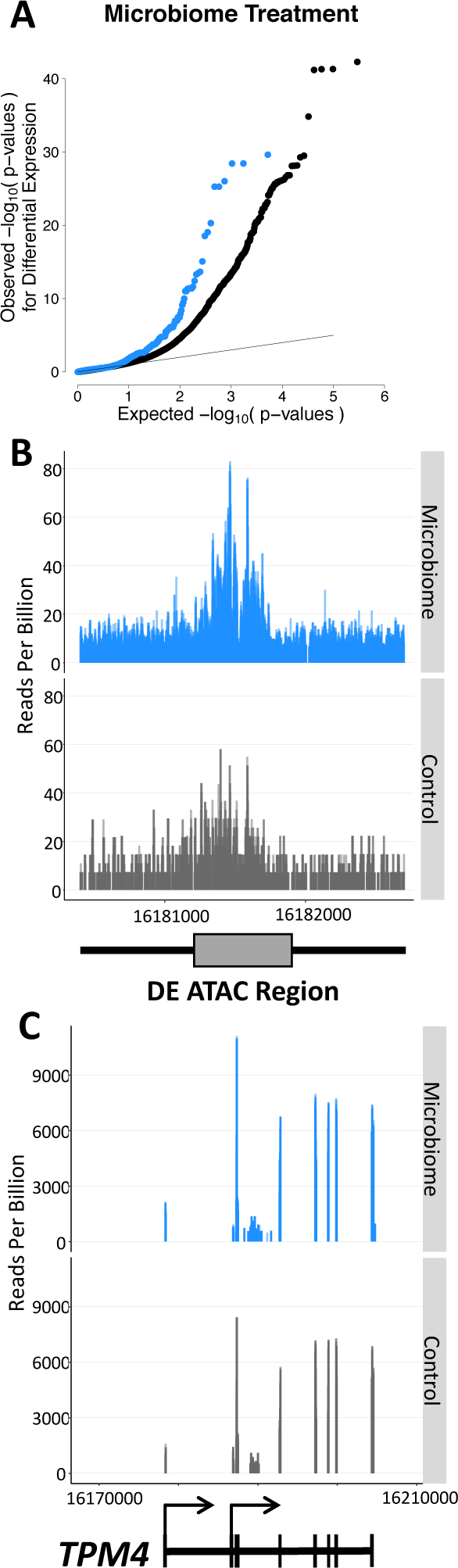
The microbiome induces changes in gene expression through changes in chromatin accessibility. **A** QQ-plot of *p*-values from the analysis of consistent differential gene expression in colonocytes treated with five microbiota samples for 4 hours. The bluepoints indicate the *p*-values for transcripts that are within 50kb of a differentially accessible region identified through ATAC-seq after 2 hours of treatment. The gray points are for transcripts not within 50kb of a differentially accessible region. **B** ATAC-seq profile centered on the 300bp window (with 1000bp on either side) that is differentially accessible following 2 hours of treatment with the microbiome (BH FDR = 12%). This region is found 4,795bp from the differentially expressed gene *TPM4*. **C** RNA-seq profile of *TPM4* (ENST00000586833) (and 20,000bp in either direction) which is differentially expressed following 4 hours of treatment with the microbiome (BH FDR = 1.9%). The gene model is shown below.

Notably, changes in chromatin accessibility and in gene expression have concordant direction, which indicates that opening of the chromatin results in up-regulation of gene expression, as expected (Fig S11). One interesting example of a gene that is regulated by the microbiome through changes in chromatin accessibility is *TPM4* which binds actin filaments and shows increased expression at 4 hours following treatment with the five microbiota samples (Fig 4B and C and Fig S12). We identified the chromatin change at a region located at 4,800 bp upstream of the transcription start site of *TPM4*. To identify transcription factors that may mediate the response to the microbiome, we performed footprinting analysis on the ATAC-seq data using CENTIPEDE [42]. We identified 397 factors active in the treatment or control conditions, for a total of 36,565,954 and 28,922,773 footprints, respectively. We identified 40 motifs with footprints in differentially accessible regions including factors that bind the ELF5, SOX2 and HMGA2 motifs (Table S12). Interestingly, when we consider a stratified FDR approach to identify differential chromatin accessibility for regions that contain footprints for active factors, we identify a larger number of differentially accessible regions (209 FDR=10%, Table S13 and Fig S10).

## Discussion

The gut microbiota has recently been associated with several different diseases and disorders [3, 7, 17, 20, 27, 50, 54]. However, the mechanism of action is not well understood, and we know little about how microbiome risk factors interact with host genes that have been linked to the same disorders. Here, we investigated how inter-individual variation in microbiome composition affects host gene regulation. Previous work in mice has shown that transplantation with microbiomes derived from mice with different phenotypes, e.g. obese versus lean, can lead to differences in organismal phenotypes [54]. In contrast, our work focuses on variable responses when considering microbiomes derived from healthy humans. Despite potential limitations due to the nature of the in vitro system used, including the presence of low level oxygen (5%) which may be deleterious to obligate anaerobes and the short duration of the treatment (up to 4hrs), we provide a first characterization of the direct effect of microbial communities and isolates on gene regulation in human colonic epithelial cells. We illustrate that while there are thousands of genes that show consistent response induced by the five microbiome treatments, there are several hundred that respond differently to each microbiome. Furthermore, some of the differences in response can be understood through the variation in microbiome composition of the five samples. Though we focus on healthy microbiomes, our results highlight the interactions of particular microbes and host genes that may play roles in various host traits. For example, we identify hundreds of genes that respond to particular microbes in a predictable way, such as the gene *GLTP* for which the response is correlated with *Collinsella*. Both *GLTP* and *Collinsella* have been associated with metabolic syndrome traits [9, 59] and our results now suggest that *Collinsella* may impact complex traits through regulation of host gene expression. While we closely examined the effect of *Collinsella* on host gene expression, there were many other relevant microbes that may also play a role in host gene regulation. For example, the abundance of *Prevotella*, which has been associated with colorectal carcinoma [51] and ileal Crohn’s disease [34], induces changes in the expression of 6 genes (BH FDR = 10%). Further studies with a wider panel of microbiome samples will allow for increased power to detect more genes that are differentially expressed across microbiome treatments.

One consideration for our analysis of the effect of *Collinsella* on host gene expression is that our treatments consist of healthy microbiota samples spiked with *Collinsella*. This was done in order to better replicate natural variation in gut microbiota composition and microbial interaction in the human gut. Microbes are constantly in competition for particular niches in the gut and so we do not expect that increases in *Collinsella* in an organism would occur alone but rather likely affect the composition of other microbes. However, future work using treatments of *Collinsella* alone may identify genes whose expression change as direct consequences of *Collinsella* abundance rather than as a result of overall changes in microbial environment induced by manipulation of *Collinsella* abundance.

Our study of chromatin accessibility changes after 2 hours of microbiome exposure in 12 samples identifies a limited number of differentially accessible regions. These results are similar to previous work in conventionalized mice that did not show broad changes in chromatin accessibility compared to germ-free mice, and could be a consequence of limited statistical power or short duration of the treatment [8, 11]. In our study, regions that are detected as significantly differentially accessible are enriched near genes that change their expression following treatment with a consistent direction of effect. Additionally, we identify specific transcription factors that are enriched in these regions and may play a key role in mediating the gene regulatory response to the microbiome but additional data and analyses will be required in the future to get a more precise description of the regulatory grammar that controls transcription factor binding, chromatin accessibility and gene expression changes. It has been shown that epigenetic changes and changes in TF binding can be induced by the gut microbiome [28, 44], and our results confirm that they may also be one of the mechanisms in human colonic epithelial cells by which the microbiota drives host gene expression.

## Conclusion

Our results suggest that specific microbes in the microbiome may be important in regulating host gene expression in the gut, and that microbes can induce changes to a large number of genes. Furthermore, our data support the hypothesis that changes in chromatin accessibility in host cells is one of the mechanisms by which the microbiome induces changes in expression of host genes that are involved in complex traits, including those that have already been associated with microbiome composition. Finally, our work suggests a molecular mechanism by which supplementing the microbiome can influence human health through changes in host cellular response. We have shown that by manipulating microbiome composition by supplementing a single microbe, we can influence the host cell regulatory response in a predictable way. This work and future research will help to determine which microbes may be most beneficial as interventional therapy to improve one’s health.

## Methods

**Extended Materials and Methods can be found in the supplement.**

### Cell culture and treatment

Experiments were conducted using primary human colonic epithelial cells (HCoEpiC, lot: 9810) which we also term, colonocytes (ScienCell 2950). Fecal microbiota was purchased from OpenBiome as live microbial suspension in 12.5% glycerol. On the day of the experient, media was removed from the colonocytes and fresh antibiotic-free media was added, followed by inoculation with microbial extract for a final microbial ratio of 10:1 microbe:colonocyte in each well. Additional wells containing only colonocytes were also cultured in the same 6-well plate to be used as controls. This experimental protocol was already described in [45].

Following 1, 2 or 4 hours, the wells were scraped on ice, pelleted and washed with cold PBS and then resuspended in lysis buffer (Dynabeads mRNA Direct Kit) and stored at −80°C until extraction of colonocyte RNA.

### *Collinsella* Spike-in Experiment

*Collinsella aerofaciens* was purchased from ATCC (cat#: 25986) and grown in Reinforced Clostridial Medium (BD Biosciences, cat#: 218081) following manufacturer’s protocol, in anaerobic conditions. We verified that we were utilizing *Collinsella aerofaciens* by extracting the DNA with the PowerSoil kit as described below. We then PCR amplified the 16S region using primers specific to *Collinsella aerofaciens* [24] (Fig S14).

Cell culturing conditions for this experiment were the same as described above. On the day of treatment, solutions were made with the microbiota sample (from individual 4) at 10:1 to the number of colonocytes and the *Collinsella aerofaciens* was spiked into the microbiota sample at 4 dilutions: 10%, 1%, 0.1% and 0.01%. Additional wells containing only colonocytes and colonocytes with the microbiota sample (0% *Collinsella aerofaciens*) were cultured as controls on the same 12-well plate. Each treatment was performed in duplicate.

Following 2 hours of culturing, the wells were scraped on ice, pelleted and washed with cold PBS, resuspended in lysis buffer (Dynabeads mRNA Direct Kit) and stored at −80°C until extraction of colonocyte RNA (as described below).

### RNA-library preparation from colonocytes

Poly-adenylated mRNAs were isolated from thawed cell lysates using the Dynabeads mRNA Direct Kit (Ambion) following the manufacturer’s instructions. RNA-seq libraries were prepared using a protocol modified from the NEBNext Ultradirectional (NEB) library preparation protocol to use Barcodes from BIOOScientific added by ligation, as described in [38]. The libraries were then pooled and sequenced on two lanes of the Illumina Next-seq 500 in the Luca/Pique laboratory using the high output kits for 75 cycles to obtain paired-end reads for an average of over 40 million total reads per sample.

### RNA sequencing and Alignment

Reads were aligned to the hg19 human reference genome using STAR [14] (https://github.com/alexdobin/STAR/releases, version STAR 2.4.0h1), and the Ensemble reference transcriptome (version 75). We further removed reads with a quality score of < 10 (equating to reads mapped to multiple locations) and removed duplicate reads using `samtools rmdup` (http://github.com/samtools/).

### Differential Gene Expression Analysis

To identify differentially expressed (DE) genes, we used `DESeq2` [31] (`R` version `3.2.1`, `DESeq2` version `1.8.1`). Gene annotations from Ensembl version 75 were used and transcripts with fewer than 20 reads total were discarded. `coverageBed` was utilized to count reads with `-s` to account for strandedness and `-split` for BED12 input. The counts were then utilized in `DESeq2` with several models to determine changes in gene expression under various conditions. A gene was considered DE if at least one of its transcripts was DE. In order to identify genes that changed at each time point following co-culturing, we used each microbiota treatment as a replicate. With this model, we identified 1,835 genes that change after 1 hour (70% of genes increase in expression), 4,099 genes after 2 hours (53% of genes increase in expression) and 1,197 genes after 4 hours (56% increase) with BH FDR < 10%, |logFC| > 0.25 (Fig S4).

In order to identify genes that were differentially expressed at a given time point after co-culturing with a specific microbiota sample we used an alternative model which allows for a different effect of each microbiome. With this model 1,131 genes changed after 1 hour with any of the 5 samples, 3,240 after 2 hours and 1,060 after 4 hours with BH FDR < 10%, |logFC| > 0.25 (Fig 1B).

We next used the likelihood ratio test that is a part of DESeq2 to compare the 2 models above in order to identify genes whose expression changes over time are determined by the individual from which the microbiota sample was taken. In this way, we identified 409 genes at BH FDR < 10%.

In order to identify components of the microbiota samples that affect gene expression we used a model that included baseline abundance of a given taxon. Baseline abundance is the number of reads (after all samples have been rarified to the sample with the lowest read count of 141,000) for a given taxon in each of the uncultured samples. Each of the time points had the same baseline abundance. This model was run for 62 taxa that had at least 141 reads (0.1% of the total reads in a sample) in at least one of the five uncultured samples. Comparing each taxon to all genes expressed in the colonocytes, we had 9,125,927 tests. We identified 588 significant comparisons (BH FDR < 10%) comprising of 46 taxa and 121 genes (Table S7).

Finally, we analyzed the validation experiment with the spike-in of *Collinsella aerofaciens*. In order to identify genes that were differentially expressed because of the *Collinsella aerofaciens*, we used the a model that included the increasing abundances of *Collinsella aerofaciens*. We identified 1,570 genes that change expression (BH FDR = 10%, Table S10, Fig S8) depending on the abundance of *Collinsella aerofaciens*.

### 16S rRNA gene sequencing and analysis of the microbiome

Half of each culturing well and the full volume of wells with microbiota samples cultured alone were used for extraction of microbial DNA using the PowerSoil kit from MO BIO Laboratories as directed, with a few modifications (found in the supplement). Microbial DNA was also extracted from the uncultured microbial samples. 16S rRNA gene amplification and sequencing was performed at the University of Minnesota Genomics Center (UMGC), as described in Burns et al. [7].

The trimmed 16S rRNA gene sequence pairs were quality filtered (q-score > 20, using QIIME 1.8.0) resulting in 1.41, 1.06, and 1.53 million high quality reads for sample replicates 1, 2, and 3, respectively [10, 40]. OTUs were picked using the closed reference algorithm against the Greengenes database (August, 2013 release) [7, 10, 12, 40]. The resulting OTU table was analyzed to determine microbial community diversity using QIIME scripts and rarefying to 141,000 reads.

We verified that the fecal samples we utilized were similar to other healthy samples by comparing the OTUs detected to the Human Microbiome Project data [22, 36]. 16S V4 OTU and HMP V1V3 OTU tables (https://www.hmpdacc.org/HMQCP/, final OTU table) were run through QIIME’s `summarize taxa.py` and consolidated at the L3 class level.

### Determining Effect of Colonocytes on Microbiota Composition

OTUs were collapsed to the genera level using scripts in QIIME 1.9.1. In total, 292 taxa were detected across all samples and treatments. After filtering on the relative abundances of each taxon, we focused on 112 taxa.

To assess how each taxon changed in response to culturing with colonocytes, we ran two linear models (including or excluding the effect of colonocytes) and compared the goodness of fit using a likelihood ratio test. The model yielded 13/112 taxa that change significantly due to treatment with a BH FDR < 10% (Table S5).

### ATAC-seq in Colonocytes Exposed to Gut Microbiota

Colonocytes were treated as described above. Each of the 5 microbiota samples were used for treatment, in replicate. Two wells of colonocytes were untreated for 2 hours under the same culturing conditions, as controls. Following 2 hours of exposure to the microbiome, cells were collected by scraping the plate on ice. Each well was counted in order to remove 50,000 cells to be used for ATAC-seq. We followed the protocol by [6] to lyse 50,000 cells and prepare ATAC-seq libraries, with the exception that we used the Illumina Nextera Index Kit (Cat #15055290) in the PCR enrichment step and the cells were not lysed with 0.1% IGEPAL CA-630 before adding the transposase to begin the ATAC-seq protocol. Individual library fragment distributions were assessed on the Agilent Bioanalyzer and pooling proportions were determined using the qPCR Kapa library quantification kit (KAPA Biosystems). Library pools were run on the Illumina NextSeq 500 Desktop sequencer in the Luca/Pique-Regi laboratory. Barcoded libraries of ATAC-seq samples were pooled and sequenced in multiple sequencing runs for 130M 38bp PE reads (max: 159M, min: 105M). One sample, derived from treatment of the colonocytes with microbiota from individual 3 was removed from later analysis due to low sequencing coverage (63M reads).

Reads were aligned to the hg19 human reference genome using HISAT2 [26] (https://github.com/, version hisat2-2.0.4), and the Ensemble reference transcriptome (hg19) with the following options:

`HISAT2 −x <genome> −1 <fastq_R1.gz> −2 <fastq_R2.gz>`

where `<genome>` represents the location of the genome file, and `<fastqs R1.gz>` and `<fastqs R2.gz>` represent the fastq files.

The multiple sequencing runs were merged for each sample using samtools (version 2.25.0). We further removed reads with a quality score of < 10 (equating to reads mapped to multiple locations) and removed duplicate reads using samtools rmdup (http://github.com/samtools/).

### Transcription factor binding footprints with CENTIPEDE

To detect which transcription factor motifs have footprints in each condition we adapted CENTIPEDE [42] to use the fragment length information contained in the ATAC-seq in the footprint model, and to jointly use in parallel the treatment and control conditions in order to ensure that the same footprint shape is used for the same motifs in both conditions.

As in CENTIPEDE, we need to start from candidate binding sites for a given motif model. For each transcription factor we scan the entire human genome (hg19) for matches to its DNA recognition motif using position weight matrix (PWM) models from TRANSFAC and JASPAR as previously described [42]. Then for each candidate location *l* we collect all the ATAC-seq fragments which are partitioned into four binds depending on the fragment length: 1) [39-99], 2) [100-139], 3) [140-179], 4) [180-250]. For each fragment, the two Tn5 insertion sites were calculated as the position 4bp after the 5’-end in the 5’ to 3’ direction. Then for each candidate motif, a matrix ***X*** was constructed to count Tn5 insertion events: each row represented a sequence match to motif in the genome (motif instance), and each column a specific cleavage site at a relative bp and orientation with respect to the motif instance. We built a matrix 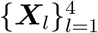 for each fragment length bin, each using a window half-size S=150bp resulting in (2 × *S* + *W*) × 2 columns, where W is the length of the motif in bp.

Finally, we fit the CENTIPEDE model in a subset of instances to determine which motifs have active binding (i.e. show footprints) with a *Z* > 5 as in [39]. The statistical significance is assessed by calculating a Z-score corresponding to the PWM effect in the prior probability in the CENTIPEDE’s logistic hierarchical prior. Then we used CENTIPEDE and motif instances with posterior probabilities higher than 0.99 to denote locations where the transcription factors are bound.

### Identification of Differentially Accessible Region Following Inoculation with Microbiota

To identify differentially accessible regions, we used `DESeq2` [31] (`R` version `3.2.1`, `DESeq2 version 1.8.1`). We separated the genome into 300bp regions and `coverageBed` was used to count reads in these regions. A total of 65,000 regions with > 0.25 reads per million were then utilized in DESeq2 using the following model:

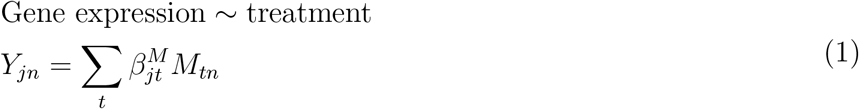

where *Y_jn_* represents the internal DEseq mean accessibility parameter for region *j* and experiment *n*, *M_n_* is the treatment indicator (control or microbiome), and 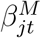 parameter is the microbiome effect. With this model, we identified 234 regions that change in the host following exposure to the gut microbiota with BH FDR < 20%. These regions were then compared to gene annotations from Ensembl version 75 to identify those that were within 50kb of a transcription start site. We found enrichment for differentially accessible regions close to DE genes at 4 hours (*p*-value = 3.96 × 10^−4^, OR = 2.13) (Fig 4A). In order to identify transcription factors which are important for response to microbiome exposure, we calculated enrichment scores for each active motif in regions of differentially accessible chromatin. 40 motifs were significantly enriched or depleted (nominal p-value < 0.05, Table S12). Because we identified factors which were enriched or depleted for differentially accessible regions, we calculated a stratified FDR for regions tested for differential chromatin accessibility by motif [52]. For each transcription factor motif, we used the footprinting results from CENTIPEDE to subset regions containing that footprint from the DESeq2 test for differential chromatin accessibility. We then adjusted the FDR using the Benjamini-Hochberg procedure [2] for each motif separately. Upon combining the results from all motifs, we identified 209 regions displaying differential chromatin accessibility with FDR < 10%.

## Supporting Information

**Supplementary File 1 — Supplemental Methods and Results**. Supplementary text for materials and methods and additional results.

## Acknowledgements

We thank members of the Luca/Pique-Regi group and Blekhman Lab for helpful discussions and comments.

## References

[1] Backhed, F., Ley, R. E., Sonnenburg, J. L., Peterson, D. A., and Gordon, J. I. (2005). Host-bacterial mutualism in the human intestine. Science, 307(5717), 1915–1920.

[2] Benjamini, Y. and Hochberg, Y. (1995). Controlling the False Discovery Rate: A Practical and Powerful Approach to Multiple Testing. Journal of the Royal Statistical Society, 57(1), 289 – 300.

[3] Bibiloni, R., Mangold, M., Madsen, K. L., Fedorak, R. N., and Tannock, G. W. (2006). The bacteriology of biopsies differs between newly diagnosed, untreated, Crohn’s disease and ulcerative colitis patients. J. Med. Microbiol., 55(Pt 8), 1141–1149.

[4] Blekhman, R., Goodrich, J. K., Huang, K., Sun, Q., Bukowski, R., Bell, J. T., Spector, T. D., Keinan, A., Ley, R. E., Gevers, D., and Clark, A. G. (2015). Host genetic variation impacts microbiome composition across human body sites. Genome Biol., 16, 191.

[5] Bolnick, D. I., Snowberg, L. K., Hirsch, P. E., Lauber, C. L., Org, E., Parks, B., Lusis, A. J., Knight, R., Caporaso, J. G., and Svanback, R. (2014). Individual diet has sex-dependent effects on vertebrate gut microbiota. Nat Commun, 5, 4500.

[6] Buenrostro, J. D., Giresi, P. G., Zaba, L. C., Chang, H. Y., and Greenleaf, W. J. (2013). Transposition of native chromatin for fast and sensitive epigenomic profiling of open chromatin, DNA-binding proteins and nucleosome position. Nat. Methods, 10(12), 1213–1218.

[7] Burns, M. B., Lynch, J., Starr, T. K., Knights, D., and Blekhman, R. (2015). Virulence genes are a signature of the microbiome in the colorectal tumor microenvironment. Genome Med, 7(1), 55.

[8] Camp, J. G., Frank, C. L., Lickwar, C. R., Guturu, H., Rube, T., Wenger, A. M., Chen, J., Bejerano, G., Crawford, G. E., and Rawls, J. F. (2014). Microbiota modulate transcription in the intestinal epithelium without remodeling the accessible chromatin landscape. Genome Res., 24(9), 1504–1516.

[9] Candela, M., Biagi, E., Soverini, M., Consolandi, C., Quercia, S., Severgnini, M., Peano, C., Turroni, S., Rampelli, S., Pozzilli, P., Pianesi, M., Fallucca, F., and Brigidi, P. (2016). Modulation of gut microbiota dysbioses in type 2 diabetic patients by macrobiotic Ma-Pi 2 diet. Br. J. Nutr., 116(1), 80–93.

[10] Caporaso, J. G., Kuczynski, J., Stombaugh, J., Bittinger, K., Bushman, F. D., Costello, E. K., Fierer, N., Pena, A. G., Goodrich, J. K., Gordon, J. I., Huttley, G. A., Kelley, S. T., Knights, D., Koenig, J. E., Ley, R. E., Lozupone, C. A., McDonald, D., Muegge, B. D., Pirrung, M., Reeder, J., Sevinsky, J. R., Turnbaugh, P. J., Walters, W. A., Widmann, J., Yatsunenko, T., Zaneveld, J., and Knight, R. (2010). QIIME allows analysis of high-throughput community sequencing data. Nat. Methods, 7(5), 335–336.

[11] Davison, J. M., Lickwar, C. R., Song, L., Breton, G., Crawford, G. E., and Rawls, J. F. (2017). Microbiota regulate intestinal epithelial gene expression by suppressing the transcription factor Hepatocyte nuclear factor 4 alpha. Genome Res., 27(7), 1195–1206.

[12] DeSantis, T. Z., Hugenholtz, P., Larsen, N., Rojas, M., Brodie, E. L., Keller, K., Huber, T., Dalevi, D., Hu, P., and Andersen, G. L. (2006). Greengenes, a chimera-checked 16S rRNA gene database and workbench compatible with ARB. Appl. Environ. Microbiol., 72(7), 5069–5072.

[13] Dethlefsen, L. and Relman, D. A. (2011). Incomplete recovery and individualized responses of the human distal gut microbiota to repeated antibiotic perturbation. Proc. Natl. Acad. Sci. U.S.A., 108 Suppl 1, 4554–4561.

[14] Dobin, A., Davis, C. A., Schlesinger, F., Drenkow, J., Zaleski, C., Jha, S., Batut, P., Chaisson, M., and Gingeras, T. R. (2013). STAR: ultrafast universal RNA-seq aligner. Bioinformatics, 29(1), 15–21.

[15] Dominianni, C., Sinha, R., Goedert, J. J., Pei, Z., Yang, L., Hayes, R. B., and Ahn, J. (2015). Sex, body mass index, and dietary fiber intake influence the human gut microbiome. PLoS ONE, 10(4), e0124599.

[16] Donohoe, D. R., Garge, N., Zhang, X., Sun, W., O’Connell, T. M., Bunger, M. K., and Bultman, S. J. (2011). The microbiome and butyrate regulate energy metabolism and autophagy in the mammalian colon. Cell Metab., 13(5), 517–526.

[17] Frank, D. N., St Amand, A. L., Feldman, R. A., Boedeker, E. C., Harpaz, N., and Pace, N. R. (2007). Molecular-phylogenetic characterization of microbial community imbalances in human inflammatory bowel diseases. Proc. Natl. Acad. Sci. U.S.A., 104(34), 13780–13785.

[18] Fujimura, K. E., Sitarik, A. R., Havstad, S., Lin, D. L., Levan, S., Fadrosh, D., Panzer, A. R., LaMere, B., Rackaityte, E., Lukacs, N. W., Wegienka, G., Boushey, H. A., Ownby, D. R., Zoratti, E. M., Levin, M., Johnson, C. C., and Lynch, S. V. (2016). Neonatal gut microbiota associates with childhood multisensitized atopy and T cell differentiation. Nat. Med., 22(10), 1187–1191.

[19] Gevers, D., Kugathasan, S., Denson, L. A., Vazquez-Baeza, Y., Van Treuren, W., Ren, B., Schwager, E., Knights, D., Song, S. J., Yassour, M., Morgan, X. C., Kostic, A. D., Luo, C., Gonzalez, A., McDonald, D., Haberman, Y., Walters, T., Baker, S., Rosh, J., Stephens, M., Heyman, M., Markowitz, J., Baldassano, R., Griffiths, A., Sylvester, F., Mack, D., Kim, S., Crandall, W., Hyams, J., Huttenhower, C., Knight, R., and Xavier, R. J. (2014). The treatment-naive microbiome in new-onset Crohn’s disease. Cell Host Microbe, 15(3), 382–392.

[20] Goodrich, J. K., Waters, J. L., Poole, A. C., Sutter, J. L., Koren, O., Blekhman, R., Beaumont, M., Van Treuren, W., Knight, R., Bell, J. T., Spector, T. D., Clark, A. G., and Ley, R. E. (2014). Human genetics shape the gut microbiome. Cell, 159(4), 789–799.

[21] Hirata, S. I. and Kunisawa, J. (2017). Gut microbiome, metabolome, and allergic diseases. Allergol Int.

[22] Huttenhower, C., Gevers, D., Knight, R., Abubucker, S., Badger, J. H., Chinwalla, A. T., Creasy, H. H., Earl, A. M., FitzGerald, M. G., Fulton, R. S., Giglio, M. G., Hallsworth-Pepin, K., Lobos, E. A., Madupu, R., Magrini, V., Martin, J. C., Mitreva, M., Muzny, D. M., Sodergren, E. J., Versalovic, J., Wollam, A. M., Worley, K. C., Wortman, J. R., Young, S. K., Zeng, Q., Aagaard, K. M., Abolude, O. O., Allen-Vercoe, E., Alm, E. J., Alvarado, L., Andersen, G. L., Anderson, S., Appelbaum, E., Arachchi, H. M., Armitage, G., Arze, C. A., Ayvaz, T., Baker, C. C., Begg, L., Belachew, T., Bhonagiri, V., Bihan, M., Blaser, M. J., Bloom, T., Bonazzi, V., Brooks, J., Buck, G. A., Buhay, C. J., Busam, D. A., Campbell, J. L., Canon, S. R., Cantarel, B. L., Chain, P. S., Chen, I. M., Chen, L., Chhibba, S., Chu, K., Ciulla, D. M., Clemente, J. C., Clifton, S. W., Conlan, S., Crabtree, J., Cutting, M. A., Davidovics, N. J., Davis, C. C., DeSantis, T. Z., Deal, C., Delehaunty, K. D., Dewhirst, F. E., Deych, E., Ding, Y., Dooling, D. J., Dugan, S. P., Dunne, W. M., Durkin, A., Edgar, R. C., Erlich, R. L., Farmer, C. N., Farrell, R. M., Faust, K., Feldgarden, M., Felix, V. M., Fisher, S., Fodor, A. A., Forney, L. J., Foster, L., Di Francesco, V., Friedman, J., Friedrich, D. C., Fronick, C. C., Fulton, L. L., Gao, H., Garcia, N., Giannoukos, G., Giblin, C., Giovanni, M. Y., Goldberg, J. M., Goll, J., Gonzalez, A., Griggs, A., Gujja, S., Haake, S. K., Haas, B. J., Hamilton, H. A., Harris, E. L., Hepburn, T. A., Herter, B., Hoffmann, D. E., Holder, M. E., Howarth, C., Huang, K. H., Huse, S. M., Izard, J., Jansson, J. K., Jiang, H., Jordan, C., Joshi, V., Katancik, J. A., Keitel, W. A., Kelley, S. T., Kells, C., King, N. B., Knights, D., Kong, H. H., Koren, O., Koren, S., Kota, K. C., Kovar, C. L., Kyrpides, N. C., La Rosa, P. S., Lee, S. L., Lemon, K. P., Lennon, N., Lewis, C. M., Lewis, L., Ley, R. E., Li, K., Liolios, K., Liu, B., Liu, Y., Lo, C. C., Lozupone, C. A., Lunsford, R., Madden, T., Mahurkar, A. A., Mannon, P. J., Mardis, E. R., Markowitz, V. M., Mavromatis, K., McCorrison, J. M., McDonald, D., McEwen, J., McGuire, A. L., McInnes, P., Mehta, T., Mihindukulasuriya, K. A., Miller, J. R., Minx, P. J., Newsham, I., Nusbaum, C., O’Laughlin, M., Orvis, J., Pagani, I., Palaniappan, K., Patel, S. M., Pearson, M., Peterson, J., Podar, M., Pohl, C., Pollard, K. S., Pop, M., Priest, M. E., Proctor, L. M., Qin, X., Raes, J., Ravel, J., Reid, J. G., Rho, M., Rhodes, R., Riehle, K. P., Rivera, M. C., Rodriguez-Mueller, B., Rogers, Y. H., Ross, M. C., Russ, C., Sanka, R. K., Sankar, P., Sathirapongsasuti, J., Schloss, J. A., Schloss, P. D., Schmidt, T. M., Scholz, M., Schriml, L., Schubert, A. M., Segata, N., Segre, J. A., Shannon, W. D., Sharp, R. R., Sharpton, T. J., Shenoy, N., Sheth, N. U., Simone, G. A., Singh, I., Smillie, C. S., Sobel, J. D., Sommer, D. D., Spicer, P., Sutton, G. G., Sykes, S. M., Tabbaa, D. G., Thiagarajan, M., Tomlinson, C. M., Torralba, M., Treangen, T. J., Truty, R. M., Vishnivetskaya, T. A., Walker, J., Wang, L., Wang, Z., Ward, D. V., Warren, W., Watson, M. A., Wellington, C., Wetterstrand, K. A., White, J. R., Wilczek-Boney, K., Wu, Y., Wylie, K. M., Wylie, T., Yandava, C., Ye, L., Ye, Y., Yooseph, S., Youmans, B. P., Zhang, L., Zhou, Y., Zhu, Y., Zoloth, L., Zucker, J. D., Birren, B. W., Gibbs, R. A., Highlander, S. K., Methe, B. A., Nelson, K. E., Petrosino, J. F., Weinstock, G. M., Wilson, R. K., and White, O. (2012). Structure, function and diversity of the healthy human microbiome. Nature, 486(7402), 207–214.

[23] Jernberg, C., Lofmark, S., Edlund, C., and Jansson, J. K. (2010). Long-term impacts of antibiotic exposure on the human intestinal microbiota. Microbiology (Reading, Engl.), 156(Pt 11), 3216–3223.

[24] Kageyama, A., Sakamoto, M., and Benno, Y. (2000). Rapid identification and quantification of Collinsella aerofaciens using PCR. FEMS Microbiol. Lett., 183(1), 43–47.

[25] Kassinen, A., Krogius-Kurikka, L., Makivuokko, H., Rinttila, T., Paulin, L., Corander, J., Malinen, E., Apajalahti, J., and Palva, A. (2007). The fecal microbiota of irritable bowel syndrome patients differs significantly from that of healthy subjects. Gastroenterology, 133(1), 24–33.

[26] Kim, D., Langmead, B., and Salzberg, S. L. (2015). HISAT: a fast spliced aligner with low memory requirements. Nat. Methods, 12(4), 357–360.

[27] Knights, D., Silverberg, M. S., Weersma, R. K., Gevers, D., Dijkstra, G., Huang, H., Tyler, A. D., van Sommeren, S., Imhann, F., Stempak, J. M., Huang, H., Vangay, P., Al-Ghalith, G. A., Russell, C., Sauk, J., Knight, J., Daly, M. J., Huttenhower, C., and Xavier, R. J. (2014). Complex host genetics influence the microbiome in inflammatory bowel disease. Genome Med, 6(12), 107.

[28] Krautkramer, K. A., Kreznar, J. H., Romano, K. A., Vivas, E. I., Barrett-Wilt, G. A., Rabaglia, M. E., Keller, M. P., Attie, A. D., Rey, F. E., and Denu, J. M. (2016). Diet-Microbiota Interactions Mediate Global Epigenetic Programming in Multiple Host Tissues. Mol. Cell, 64(5), 982–992.

[29] Larsen, N., Vogensen, F. K., van den Berg, F. W., Nielsen, D. S., Andreasen, A. S., Pedersen, B. K., Al-Soud, W. A., Sørensen, S. J., Hansen, L. H., and Jakobsen, M. (2010). Gut microbiota in human adults with type 2 diabetes differs from non-diabetic adults. PLoS ONE, 5(2), e9085.

[30] Lee, S., Sung, J., Lee, J., and Ko, G. (2011). Comparison of the gut microbiotas of healthy adult twins living in South Korea and the United States. Appl. Environ. Microbiol., 77(20), 7433–7437.

[31] Love, M. I., Huber, W., and Anders, S. (2014). Moderated estimation of fold change and dispersion for RNA-seq data with DESeq2. Genome Biol., 15(12), 550.

[32] Lozupone, C. A., Stombaugh, J. I., Gordon, J. I., Jansson, J. K., and Knight, R. (2012). Diversity, stability and resilience of the human gut microbiota. Nature, 489(7415), 220–230.

[33] Makivuokko, H., Tiihonen, K., Tynkkynen, S., Paulin, L., and Rautonen, N. (2010). The effect of age and non-steroidal anti-inflammatory drugs on human intestinal microbiota composition. Br. J. Nutr., 103(2), 227–234.

[34] Manichanh, C., Rigottier-Gois, L., Bonnaud, E., Gloux, K., Pelletier, E., Frangeul, L., Nalin, R., Jarrin, C., Chardon, P., Marteau, P., Roca, J., and Dore, J. (2006). Reduced diversity of faecal microbiota in Crohn’s disease revealed by a metagenomic approach. Gut, 55(2), 205–211.

[35] Maurice, C. F., Haiser, H. J., and Turnbaugh, P. J. (2013). Xenobiotics shape the physiology and gene expression of the active human gut microbiome. Cell, 152(1-2), 39–50.

[36] Methe, B. A., Nelson, K. E., Pop, M., Creasy, H. H., Giglio, M. G., Huttenhower, C., Gevers, D., Petrosino, J. F., Abubucker, S., Badger, J. H., Chinwalla, A. T., Earl, A. M., FitzGerald, M. G., Fulton, R. S., Hallsworth-Pepin, K., Lobos, E. A., Madupu, R., Magrini, V., Martin, J. C., Mitreva, M., Muzny, D. M., Sodergren, E. J., Versalovic, J., Wollam, A. M., Worley, K. C., Wortman, J. R., Young, S. K., Zeng, Q., Aagaard, K. M., Abolude, O. O., Allen-Vercoe, E., Alm, E. J., Alvarado, L., Andersen, G. L., Anderson, S., Appelbaum, E., Arachchi, H. M., Armitage, G., Arze, C. A., Ayvaz, T., Baker, C. C., Begg, L., Belachew, T., Bhonagiri, V., Bihan, M., Blaser, M. J., Bloom, T., Bonazzi, V. R., Brooks, P., Buck, G. A., Buhay, C. J., Busam, D. A., Campbell, J. L., Canon, S. R., Cantarel, B. L., Chain, P. S., Chen, I. M., Chen, L., Chhibba, S., Chu, K., Ciulla, D. M., Clemente, J. C., Clifton, S. W., Conlan, S., Crabtree, J., Cutting, M. A., Davidovics, N. J., Davis, C. C., DeSantis, T. Z., Deal, C., Delehaunty, K. D., Dewhirst, F. E., Deych, E., Ding, Y., Dooling, D. J., Dugan, S. P., Dunne, W., Durkin, A., Edgar, R. C., Erlich, R. L., Farmer, C. N., Farrell, R. M., Faust, K., Feldgarden, M., Felix, V. M., Fisher, S., Fodor, A. A., Forney, L., Foster, L., Di Francesco, V., Friedman, J., Friedrich, C., Fronick, C. C., Fulton, L. L., Gao, H., Garcia, N., Giannoukos, G., Giblin, C., Giovanni, M. Y., Goldberg, J. M., Goll, J., Gonzalez, A., Griggs, A., Gujja, S., Haas, B. J., Hamilton, H. A., Harris, L., Hepburn, T. A., Herter, B., Hoffmann, D. E., Holder, M. E., Howarth, C., Huang, K. H., Huse, S. M., Izard, J., Jansson, J. K., Jiang, H., Jordan, C., Joshi, V., Katancik, J. A., Keitel, W. A., Kelley, S. T., Kells, C., Kinder-Haake, S., King, N. B., Knight, R., Knights, D., Kong, H. H., Koren, O., Koren, S., Kota, K. C., Kovar, C. L., Kyrpides, N. C., La Rosa, P. S., Lee, S. L., Lemon, K. P., Lennon, N., Lewis, C. M., Lewis, L., Ley, R. E., Li, K., Liolios, K., Liu, B., Liu, Y., Lo, C. C., Lozupone, C. A., Lunsford, R., Madden, T., Mahurkar, A. A., Mannon, P. J., Mardis, E. R., Markowitz, V. M., Mavrommatis, K., McCorrison, J. M., McDonald, D., McEwen, J., McGuire, A. L., McInnes, P., Mehta, T., Mihindukulasuriya, K. A., Miller, J. R., Minx, P. J., Newsham, I., Nusbaum, C., O’Laughlin, M., Orvis, J., Pagani, I., Palaniappan, K., Patel, S. M., Pearson, M., Peterson, J., Podar, M., Pohl, C., Pollard, K. S., Priest, M. E., Proctor, L. M., Qin, X., Raes, J., Ravel, J., Reid, J. G., Rho, M., Rhodes, R., Riehle, K. P., Rivera, M. C., Rodriguez-Mueller, B., Rogers, Y. H., Ross, M. C., Russ, C., Sanka, R. K., Sankar, P., Sathirapongsasuti, J., Schloss, J. A., Schloss, P. D., Schmidt, T. M., Scholz, M., Schriml, L., Schubert, A. M., Segata, N., Segre, J. A., Shannon, W. D., Sharp, R. R., Sharpton, T. J., Shenoy, N., Sheth, N. U., Simone, G. A., Singh, I., Smillie, C. S., Sobel, J. D., Sommer, D. D., Spicer, P., Sutton, G. G., Sykes, S. M., Tabbaa, D. G., Thiagarajan, M., Tomlinson, C. M., Torralba, M., Treangen, T. J., Truty, R. M., Vishnivetskaya, T. A., Walker, J., Wang, L., Wang, Z., Ward, D. V., Warren, W., Watson, M. A., Wellington, C., Wetterstrand, K. A., White, J. R., Wilczek-Boney, K., Wu, Y. Q., Wylie, K. M., Wylie, T., Yandava, C., Ye, L., Ye, Y., Yooseph, S., Youmans, B. P., Zhang, L., Zhou, Y., Zhu, Y., Zoloth, L., Zucker, J. D., Birren, B. W., Gibbs, R. A., Highlander, S. K., Weinstock, G. M., Wilson, R. K., and White, O. (2012). A framework for human microbiome research. Nature, 486(7402), 215–221.

[37] Moore, W. E. and Moore, L. H. (1995). Intestinal floras of populations that have a high risk of colon cancer. Appl. Environ. Microbiol., 61(9), 3202–3207.

[38] Moyerbrailean, G. A., Davis, G. O., Harvey, C. T., Watza, D., Wen, X., Pique-Regi, R., and Luca, F. (2015). A high-throughput RNA-seq approach to profile transcriptional responses. Sci Rep, 5, 14976.

[39] Moyerbrailean, G. A., Kalita, C. A., Harvey, C. T., Wen, X., Luca, F., and Pique-Regi, R. (2016). Which Genetics Variants in DNase-Seq Footprints Are More Likely to Alter Binding? PLoS Genet., 12(2), e1005875.

[40] Navas-Molina, J. A., Peralta-Sanchez, J. M., Gonzalez, A., McMurdie, P. J., Vazquez-Baeza, Y., Xu, Z., Ursell, L. K., Lauber, C., Zhou, H., Song, S. J., Huntley, J., Ackermann, G. L., Berg-Lyons, D., Holmes, S., Caporaso, J. G., and Knight, R. (2013). Advancing our understanding of the human microbiome using QIIME. Meth. Enzymol., 531, 371–444.

[41] Ott, S. J., Musfeldt, M., Wenderoth, D. F., Hampe, J., Brant, O., Folsch, U. R., Timmis, K. N., and Schreiber, S. (2004). Reduction in diversity of the colonic mucosa associated bacterial microflora in patients with active inflammatory bowel disease. Gut, 53(5), 685–693.

[42] Pique-Regi, R., Degner, J. F., Pai, A. A., Gaffney, D. J., Gilad, Y., and Pritchard, J. K. (2011). Accurate inference of transcription factor binding from DNA sequence and chromatin accessibility data. Genome Res., 21(3), 447–455.

[43] Qin, J., Li, R., Raes, J., Arumugam, M., Burgdorf, K. S., Manichanh, C., Nielsen, T., Pons, N., Levenez, F., Yamada, T., Mende, D. R., Li, J., Xu, J., Li, S., Li, D., Cao, J., Wang, B., Liang, H., Zheng, H., Xie, Y., Tap, J., Lepage, P., Bertalan, M., Batto, J. M., Hansen, T., Le Paslier, D., Linneberg, A., Nielsen, H. B., Pelletier, E., Renault, P., Sicheritz-Ponten, T., Turner, K., Zhu, H., Yu, C., Li, S., Jian, M., Zhou, Y., Li, Y., Zhang, X., Li, S., Qin, N., Yang, H., Wang, J., Brunak, S., Dore, J., Guarner, F., Kristiansen, K., Pedersen, O., Parkhill, J., Weissenbach, J., Bork, P., Ehrlich, S. D., Wang, J., Antolin, M., Artiguenave, F., Blottiere, H., Borruel, N., Bruls, T., Casellas, F., Chervaux, C., Cultrone, A., Delorme, C., Denariaz, G., Dervyn, R., Forte, M., Friss, C., van de Guchte, M., Guedon, E., Haimet, F., Jamet, A., Juste, C., Kaci, G., Kleerebezem, M., Knol, J., Kristensen, M., Layec, S., Le Roux, K., Leclerc, M., Maguin, E., Minardi, R. M., Oozeer, R., Rescigno, M., Sanchez, N., Tims, S., Torrejon, T., Varela, E., de Vos, W., Winogradsky, Y., and Zoetendal, E. (2010). A human gut microbial gene catalogue established by metagenomic sequencing. Nature, 464(7285), 59–65.

[44] Qin, Y., Roberts, J. D., Grimm, S. A., Lih, F. B., Deterding, L. J., Li, R., Chrysovergis, K., and Wade, P. A. (2018). An obesity-associated gut microbiome reprograms the intestinal epigenome and leads to altered colonic gene expression. Genome Biol., 19(1), 7.

[45] Richards, A. L., Burns, M. B., Alazizi, A., Barreiro, L. B., Pique-Regi, R., Blekhman, R., and Luca, F. (2016). Genetic and transcriptional analysis of human host response to healthy gut microbiota. mSystems, 1(4).

[46] Rowland, I., Gibson, G., Heinken, A., Scott, K., Swann, J., Thiele, I., and Tuohy, K. (2017). Gut microbiota functions: metabolism of nutrients and other food components. Eur J Nutr.

[47] Schwiertz, A., Taras, D., Schafer, K., Beijer, S., Bos, N. A., Donus, C., and Hardt, P. D. (2010). Microbiota and SCFA in lean and overweight healthy subjects. Obesity (Silver Spring), 18(1), 190–195.

[48] Semenkovich, N. P., Planer, J. D., Ahern, P. P., Griffin, N. W., Lin, C. Y., and Gordon, J. I. (2016). Impact of the gut microbiota on enhancer accessibility in gut intraepithelial lymphocytes. Proc. Natl. Acad. Sci. U.S.A., 113(51), 14805–14810.

[49] Seto, C. T., Jeraldo, P., Orenstein, R., Chia, N., and DiBaise, J. K. (2014). Prolonged use of a proton pump inhibitor reduces microbial diversity: implications for Clostridium difficile susceptibility. Microbiome, 2, 42.

[50] Sinha, R., Ahn, J., Sampson, J. N., Shi, J., Yu, G., Xiong, X., Hayes, R. B., and Goedert, J. J. (2016). Fecal Microbiota, Fecal Metabolome, and Colorectal Cancer Interrelations. PLoS ONE, 11(3), e0152126.

[51] Sobhani, I., Tap, J., Roudot-Thoraval, F., Roperch, J. P., Letulle, S., Langella, P., Corthier, G., Tran Van Nhieu, J., and Furet, J. P. (2011). Microbial dysbiosis in colorectal cancer (CRC) patients. PLoS ONE, 6(1), e16393.

[52] Sun, L., Craiu, R. V., Paterson, A. D., and Bull, S. B. (2006). Stratified false discovery control for large-scale hypothesis testing with application to genome-wide association studies. Genet. Epidemiol., 30(6), 519–530.

[53] Tims, S., Derom, C., Jonkers, D. M., Vlietinck, R., Saris, W. H., Kleerebezem, M., de Vos, W. M., and Zoetendal, E. G. (2013). Microbiota conservation and BMI signatures in adult monozygotic twins. ISME J, 7(4), 707–717.

[54] Turnbaugh, P. J., Backhed, F., Fulton, L., and Gordon, J. I. (2008). Diet-induced obesity is linked to marked but reversible alterations in the mouse distal gut microbiome. Cell Host Microbe, 3(4), 213–223.

[55] Turnbaugh, P. J., Hamady, M., Yatsunenko, T., Cantarel, B. L., Duncan, A., Ley, R. E., Sogin, M. L., Jones, W. J., Roe, B. A., Affourtit, J. P., Egholm, M., Henrissat, B., Heath, A. C., Knight, R., and Gordon, J. I. (2009a). A core gut microbiome in obese and lean twins. Nature, 457(7228), 480–484.

[56] Turnbaugh, P. J., Ridaura, V. K., Faith, J. J., Rey, F. E., Knight, R., and Gordon, J. I. (2009b). The effect of diet on the human gut microbiome: a metagenomic analysis in humanized gnotobiotic mice. Sci Transl Med, 1(6), 6ra14.

[57] Welter, D., MacArthur, J., Morales, J., Burdett, T., Hall, P., Junkins, H., Klemm, A., Flicek, P., Manolio, T., Hindorff, L., and Parkinson, H. (2014). The NHGRI GWAS Catalog, a curated resource of SNP-trait associations. Nucleic Acids Res., 42(Database issue), D1001–1006.

[58] Yatsunenko, T., Rey, F. E., Manary, M. J., Trehan, I., Dominguez-Bello, M. G., Contreras, M., Magris, M., Hidalgo, G., Baldassano, R. N., Anokhin, A. P., Heath, A. C., Warner, B., Reeder, J., Kuczynski, J., Caporaso, J. G., Lozupone, C. A., Lauber, C., Clemente, J. C., Knights, D., Knight, R., and Gordon, J. I. (2012). Human gut microbiome viewed across age and geography. Nature, 486(7402), 222–227.

[59] Zabaneh, D. and Balding, D. J. (2010). A genome-wide association study of the metabolic syndrome in Indian Asian men. PLoS ONE, 5(8), e11961.

